# Task-Dependent and Cell-Type-Specific Modulation of the Mouse Dorsal Cortex of the Inferior Colliculus

**DOI:** 10.64898/2026.07.23.740270

**Authors:** Gang Xiao, Brandon Li, Sai Smriti Gogineni, Daniel Adolfo Llano

**Affiliations:** Department of Molecular and Integrative Physiology, University of Illinois, Urbana-Champaign, IL, 61801; The Beckman Institute for Advanced Science and Technology, University of Illinois, Urbana-Champaign, IL, 61801

## Abstract

Sensory responses in the auditory system are modulated by behavioral context, and this modulation extends as early as the inferior colliculus (IC). Where within the IC it occurs, which neurons it affects, what behavioral variables drive it, and whether it requires the auditory cortex remain unclear. We imaged the dorsal cortex of the IC (DCIC) with two-photon microscopy in head-fixed mice (n = 7) performing a two-alternative forced-choice tone-discrimination task, with blocks of passive listening to the same tones before and after, while recording orofacial movement, licking, and pupil diameter from synchronized video. Unsupervised clustering of sound-evoked responses in 1479 tracked neurons identified four functional response classes: onset-excited, sustained-inhibited, offset/motor-excited, and transient-inhibited. Task engagement modulated these classes in distinct and outcome-dependent ways, with the sustained-inhibited and transient-inhibited classes more suppressed and the offset/motor-excited class more excited during active trials, and with hit and miss trials driving most of the effect. Population activity discriminated active from passive trials above chance, and this discrimination outlasted the stimulus. A linear encoding model attributed a substantial share of the explained variance in three of the four classes to facial movement, whereas pupil diameter contributed little in any class. The correlation between orofacial motion and activity was itself context-dependent in the sustained-inhibited and transient-inhibited classes, but not in the offset/motor-excited class. Bilateral chemogenetic silencing of the auditory cortex (n = 3 mice, 171 matched neurons) altered task modulation only in the sustained-inhibited class. Task-related modulation in the DCIC is therefore heterogeneous across functionally defined neurons, is closely tied to movement, and depends on the auditory cortex in only one of these populations.

## Introduction

Sensory perception can vary depending on the behavioral context. Whether an animal engages with a stimulus actively or passively can result in a different percept, and neural activity throughout the mammalian brain is modified during task engagement in many sensory modalities, including the auditory system (Busse et al., 2017). In the auditory system this modulation has been characterized most extensively in the thalamus and cortex (Kuchibhotla et al., 2017; Yao et al., 2019), but task engagement also modulates activity as early as the inferior colliculus (IC) (De Franceschi & Barkat, 2021). Since the IC is a relatively early station in the auditory pathway, this finding suggests that perception modulation may begin much earlier than previously thought. It also leaves several questions open: where within the IC this modulation arises and which neurons it affects, what behavioral variable actually drives it, and whether it requires the cortex.

Task modulation need not be distributed uniformly within the IC. Unlike the lemniscal central nucleus of the IC (CNIC), which descending corticocollicular (CC) axons innervate only sparsely, the dorsal cortex of the IC (DCIC) is a principal target of these projections, which arise mainly from layer 5 of auditory cortex and to a smaller extent from layer 6 (Bajo & King, 2012; Schofield, 2010; Stebbings et al., 2014). The DCIC also receives neuromodulatory and non-auditory afferents that largely bypass the CNIC (Hormigo et al., 2012; Motts & Schofield, 2009; Patel et al., 2017). It is therefore positioned to receive more top-down and nonlemniscal input than the CNIC, and if task-related modulation in the IC reflects descending or non-auditory influences, the DCIC is where it should be most apparent. Modulation need not be uniform within the DCIC either. Brain state already modulates DCIC sound responses differently for excitatory and inhibitory neurons (Chen & Song, 2019). Whether task modulation is similarly heterogeneous, and how it is distributed across neurons with different sound-response profiles, is unknown. Addressing this requires classifying DCIC neurons by their responses before asking how each class is modulated.

Task engagement is also not a single variable. An animal performing a task is more aroused and moving more than the same animal listening passively, and both arousal and movement influence activity in the IC. Task engagement and pupil-indexed arousal make separable contributions to auditory responses along the pathway (Saderi et al., 2021), and locomotion-driven activity in the IC persists in completely deafened mice, indicating that movement-related activity there does not arise from self-generated sound (Han et al., 2026). An active-passive difference in DCIC activity could therefore reflect task engagement, arousal, movement, or some combination of them. Separating these possibilities requires recording arousal and movement continuously and synchronously with neural activity during the task itself, rather than treating the active-passive difference as engagement by default.

Descending input from the auditory cortex is one candidate source of task modulation in the IC. Stimulation of auditory cortex shifts the frequency, intensity, and duration tuning of IC neurons (Suga, 2020), and pathway-specific manipulations show that CC projections shape IC sound coding in ways that depend on the stimulus (Blackwell et al., 2020; Lesicko et al., 2022). Layer 5 and layer 6 CC neurons also differ in their inputs and subcortical targets (Issa et al., 2023; Slater et al., 2019), so the pathway is not a single functional channel. The pathway-specific manipulations above were carried out during passive listening, and whether the auditory cortex is required for task-dependent modulation in the DCIC has not been tested.

We developed a 2-alternative-forced-choice (2AFC) tone-discrimination paradigm conducted while imaging the DCIC through a cranial window using multiphoton microscopy, with blocks of passive presentation of the same tones before and after, and recorded orofacial movement, licking, and pupil diameter from synchronized video throughout. We investigated: 1) whether task-related modulation is present in the DCIC and whether it is uniform across functionally distinct neurons; 2) whether this modulation can be explained by behavioral variables during the task, such as movement, lick choice, and pupil size; and 3) whether it depends on the auditory cortex, tested by chemogenetic silencing. We found four functional response classes modulated by behavioral context in distinct, outcome-dependent ways; movement accounted for a substantial share of the explained variance in three of them, while pupil size contributed little in any; and silencing the auditory cortex altered the modulation of only one of these classes.

## Materials and methods

### Animals

F1 progeny from crosses between transgenic C57BL/6J mice and B6.CAST-Cdh23 Ahl+ /Kjn mice were used. Seven mice of either sex, aged 4–12 months, were included in this study. All animal procedures were approved by the Institutional Animal Care and Use Committee at the University of Illinois at Urbana-Champaign.

### Behavioral training

A 2AFC task was used as the behavioral test. A custom-made behavioral apparatus was used, and the awake mouse was head-fixed by sliding the headpost glued on the mouse’s head under the screws standing on two holding arms and tightening the screws to lock the headpost. There were two tubes near the mouth of the mouse for licking-choice detection and water reward delivery. The mice were water-restricted, so they were motivated to perform several hundred trials over the course of a training session. After habituation and learning of the tube’s location, the mice were trained to lick the specific tube according to the sound frequency they heard. In each trial, there was a pure tone presented for 500 ms at 70 dB SPL. Then, the animal was given a 2.5 s response window to respond by licking one of the two tubes (Fig. 1A). The lick choice of the mouse was only counted after the offset of the tone. Two pure tones were correlated with the two lick tubes. If they licked the tube corresponding to the sound, 5 μL sucrose water was delivered through that tube. If they licked the other tube, there was a 10 s timeout as punishment. The timeout was not explicitly cued; reward was withheld and the trial simply extended by the timeout period. The trial ended either after the response window or the time-out period. Between each trial, there was an 8-10s random inter-trial interval to prevent the prediction of the trial onset of the next trial. Accuracy was calculated as:

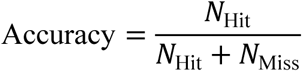

**Fig. 1.**
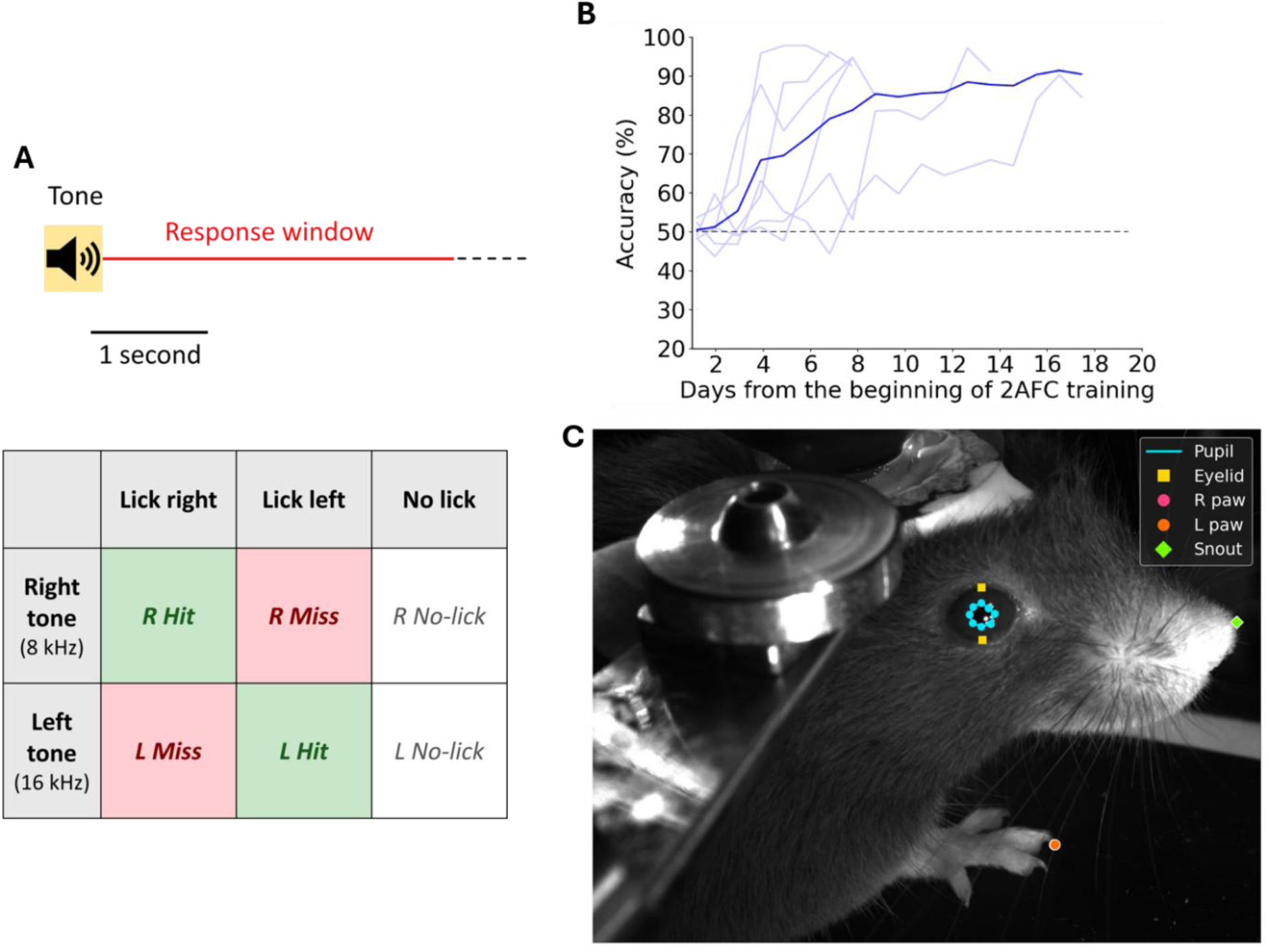
Two-alternative forced-choice tone discrimination task and behavioral learning. (A) Schematic of the 2AFC tone discrimination paradigm. Mice were presented with either an 8 kHz right target tone or a 16 kHz left target tone and were required to report the perceived tone by licking the corresponding lick port within the response window. The tone was 500 ms and the responding window was 2.5 s after the tone offset. Lick choice was only counted after the offset of the tone. For 8 kHz trials, right licks were classified as right-hit trials, left licks as right-miss trials, and trials without a lick response as right-no-lick trials. For 16 kHz trials, left licks were classified as left-hit trials, right licks as left-miss trials, and trials without a lick response as left-no-lick trials. (B) Behavioral learning curve across days from the beginning of 2AFC training. Light blue traces represent the performance of individual mice, and the dark blue trace represents the mean accuracy across mice. The dashed horizontal line indicates chance performance at 50%. Mice gradually improved their task performance over training and reached high accuracy after several days of training. Values are shown as mean accuracy of the left accuracy and right accuracy. (C) Example video frame showing DeepLabCut-based tracking of behavioral features during the task. Labeled points indicate the tracked pupil, eyelid, right paw, left paw, and snout positions. These tracking labels were used to monitor task-related behavioral variables, including pupil dynamics and orofacial/body movements, during imaging sessions.

After their accuracy was beyond 75% for both sides’ choices, the mice were ready for doing the task while 2P imaging the IC neurons. All behavioral progress was monitored and controlled by a custom MATLAB program controlling the RX6 TDT processor and several Arduino microcontrollers.

In the 2P imaging session, the mouse conducted a normal 2AFC for 100 trials. The mouse also passively listened to 100 trials with the lick tubes withdrawn and without the task and rewards. 50 passive trials took place before the behavioral trials and 50 after.

### Behavioral video recording

During imaging, the mouse’s face and body were recorded using two cameras: a face camera (acA1440-220um, Basler) positioned to capture orofacial and licking movements as well as the pupil, and a back camera (acA800-510um, Basler) capturing body movement. Both cameras imaged under infrared illumination (850 nm), allowing the mouse to be recorded in darkness without introducing visible light that could affect the 2P microscope. Because the pupil dilates and approaches saturation under these dark conditions, leaving little measurable dynamic range, a UVA light source (365 nm) was directed at the eye contralateral to the recorded pupil. This evoked a consensual pupillary constriction that held the recorded pupil at a mid-range diameter, preserving its dynamics for measurement. UVA was used because its wavelength lies outside the IR imaging and 2P detection ranges and therefore does not interfere with image acquisition. Both cameras and the 2P imaging system were triggered by TTL pulses from the TDT processor (RX6, TDT), which also controlled the behavioral task and acoustic stimuli, ensuring that video frames, 2P frames, and behavioral events (sound onset and licking) were synchronized to a common time base. The video was recorded in 40 fps.

### Motion energy

Motion energy (ME) was computed as the mean absolute pixel-intensity difference between consecutive video frames of the face camera. Whole facial ME (movement) was computed over the entire frame with the eye region excluded, to prevent pupil and eye movements from contributing to the movement signal. Mouth ME was computed identically but restricted to a manually defined region of interest around the mouth, capturing orofacial and licking movements.

### Craniotomy surgery

Before surgery, mice were anesthetized with isoflurane (4% induction, 1–2% maintenance) and placed into a stereotaxic frame (Kopf Model 940). To prevent brain edema during or after the craniotomy, an intramuscular injection of dexamethasone sodium (4.8 mg/kg) was given just before the surgery. After fixation, both eyes of the mouse were protected by applying Opti care lubricant eye gel (Aventix Animal Health). The hair on the scalp was then removed by massaging the scalp with a depilatory cream (Nair) and leaving the cream on the scalp for 3 min. The cream was then removed by a thin plastic sheet (flexible ruler) to leave a hair-free area on the scalp. The remaining tiny hairs were then removed by saline swab and the area was then sterilized by applying 10% povidone iodine (Dynarex). The medial incision was made with a scalpel blade #10, and 0.2 ml of 0.5% lidocaine was injected intradermally into the scalp. The skin extending from the medial line to the temporalis muscle was completely removed using a pair of microscissors to produce a wide skinless area above the skull. A pair of no. 5/45 forceps were used to remove any remaining periosteum. The skull was cleaned with sterile saline and dried with gentle pressurized air. The bone surface was slightly etched by dental-grade 35% phosphoric acid gel for 20 seconds. A 3.5 mm diameter circle was drawn on the skull above the left IC. A handheld drill (Foredom) with 0.2 mm burr drill bit was used to drill through the skull.

To prevent overheating of the superficial brain tissues and to mitigate the occasional spurts of skull bleeding during the drilling, ice-cold sterile saline was used to intermittently irrigate the surface. A stream of pressurized air was also applied during the drilling procedure to prevent overheating and remove the debris produced by the drilling. Caution was taken while crossing the sagittal or the lambdoid sutures to avoid damaging the underlying sinuses. After drilling, the skull was irrigated in sterile saline and the bone flap (the undrilled bone over the craniotomy area) was gently removed using a pair of no. 5/45 forceps. To control the bleeding if it occurred, a piece of sterile hemostatic gel which was presoaked in ice-cold saline, was applied to the bleeding spot. Once the bleeding ceased, the brain was kept covered in sterile saline. In some surgeries, the dura needed to be peeled off. A 30 G needle tip was used to pierce the dura in the medial area between the two ICs and devoid of cortical vasculature. After piercing, the dura was carefully and gently lifted, and a Bone microprobe was used to grab the dura to gently tear it to the extent of the transverse sinus to avoid bleeding. AAV-hSyn-Soma-jGCaMP8m virus was injected into 3 sites x 2 depth in the DCIC with each injection volume of 25 nL. A 3D-printed biocompatible plastic well was glued to a 3 mm diameter glass coverslip with NOA 68 glue as the cover glass. The cover glass was secured by the drill bit by gently pressing the cover glass from the top. The cover glass was further pressed down to plug and hold the brain in place. The skull was covered with C&B Metabond dental cement to make a base and glue the cover glass. A custom-made titanium head post was glued carefully on the top of the skull to be at the same level as the cover glass also with the dental cement.

### 2-photon imaging

A custom-built 2P microscope was used. The optical and the controlling components were supplied from Bruker, Olympus, and Thorlabs. The imaging of the DCIC was made using a 16× water-immersion objective (N16XLWD-PF, 16×, NA: 0.80, WD: 3 mm; Nikon). For imaging both the GCaMP8m signals, the excitation light was generated by InSight X3 laser (Spectra-Physics Lasers) tuned to a wavelength of 920 or 1040 nm, respectively. A layer of diluted multipurpose wavelengths ultrasound gel (National therapy) with deionized water was used to immerse the objective. This gel was able to trap the water and reduce its evaporation during imaging. The emitted signals were detected by a photomultiplier tube (Hamamatsu H7422PA-4) following a t565lp dichroic and a Chroma barrier et525/70m filter for GCaMP8m. Images (512 × 512 pixels) were collected at a frame rate of 29.5 Hz in the resonant galvo mode. The field of view was selected based on the expression of GCaMP8m and being acoustically active using a search stimulus that was 500-ms broadband noise with zero modulation at 80 dB SPL. The imaging timing was controlled with digital signals sent from the TDT processors. The TDT processors were controlled by the same custom MATLAB program which controlled the behavior process to synchronize the two.

The imaging was accompanied by the mouse’s active and passive listening behaviors. In every trial of the behavior session, 2 s before and 2 s after the sound onset were imaged.

In experiments with AC silencing, mice were injected intraperitoneally with 3 mg/kg CNO. In the control days, they were injected with the same amount of saline.

### Analysis of 2P images

The collected 2P frame stacks were motion corrected by the Python package suite2p. ROIs of neurons were manually drawn on the averaged image of the motion-corrected frame stack according to the morphology of the neuron (dark nucleus with bright rim). The matched ROIs were marked in the study with imaging happening on different days. The ROIs were then used to create two masks; one mask was used as a replica of the ROIs and the second mask was made around the original ROI (roughly four times larger in the area). The smaller mask was applied to find the average pixel value within each ROI, while the larger mask was applied to find the average pixel value of the neuropil. The neuropil correction was applied using the following equation (Akerboom et al., 2012):

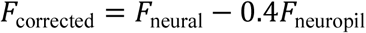

ΔF/F values were then calculated by using the following equation.

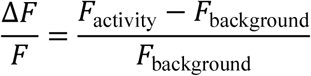

where the background value is the mean value of masks between 0.5-0 s prior to the sound onset.

The modulation index was calculated by:

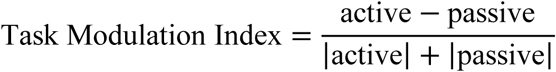

### Stereotaxic injection surgery

Mice were anesthetized with isoflurane (4% induction, 1–2% maintenance) and placed into a stereotaxic frame (Kopf Model 940). For validation of the AC silencing with chemogenetic method, pressure injections of *Cre*-dependent pGP-AAV-syn-FLEX-jGCaMP8m-WPRE and CAV hSyn DIO hM4D-mCherry were injected at 2 sites in the left AC of the Rbp4cre mice (1.3 and 1.7 mm anterior to lambdoid suture at the temporal ridge) at depths of 600–800 μm and an angle of 40 degrees from the sagittal plane. For bilateral AC silencing with chemogenetic method, pressure injections of pAAV-CaMKIIa-hM4D(Gi)-mCherry (AAV2) were injected at 2 sites in the bilateral AC (1.3 and 1.7 mm anterior to lambdoid suture at the temporal ridge) at depths of 700–1000 μm and an angle of 40 degrees from the sagittal plane.

### Unsupervised clustering

Responsive DCIC neurons were identified by a Benjamini–Hochberg-corrected Wilcoxon signed-rank test (q < 0.05, two-sided) comparing the sound (0–0.5 s) or post-sound (0.5–1.5 s) window to the pre-sound baseline (−0.5–0 s) across all trials. For each neuron, the mean PSTH across active trials with the same pure tone frequency was z-scored across time (−1 to 2 s). Consensus k-means clustering (k = 3, 200 runs, first 3 PCs) followed by average-linkage hierarchical clustering of the co-association matrix yielded three base clusters. Onset-excited neurons were extracted post-hoc using a one-sided signed-rank test on per-trial sound window (0–0.5 s) vs. baseline responses (BH q < 0.05), positive z-scored mean, and response ΔF/F > 0.3, giving four final clusters.

For each functional cluster, mean ΔF/F was computed separately for active trials (or hit\miss\no-lick trials separately) and passive trials, baseline-subtracted using the −0.5–0 s pre-tone window. Responses were quantified in two windows: the sound window (0–0.5 s) and the post-sound window (0.5–1.5 s), which was the mean of the ΔF/F in the window.

### Support vector machine

To test whether DCIC population activity carries context-dependent information beyond sensory responses, we performed time-resolved linear decoding of behavioral context (active vs. passive) using the same tone stimulus trials. At each time point, the instantaneous ΔF/F of all simultaneously recorded neurons served as features for a linear SVM classifier trained to discriminate active from passive trials. Classification accuracy was estimated using stratified 5-fold cross-validation within each session (n = 7 sessions). Decoding time courses were averaged across sessions and are shown as mean ± SEM.

### Neural encoding model

To decompose each neuron’s activity variance into contributions from sensory, motor, and arousal signals, we fit a linear ridge-regression encoding model to single-trial ΔF/F traces. Only active trials were included. Four predictor groups were constructed:

#### Sound identity

Three binary regressors (8 kHz, 16 kHz, 11314 Hz (an ambiguous tone presented in some active trials)) set to 1 during the sound window (0–0.5 s) and convolved with an exponential calcium indicator kernel (τ = 0.5 s) to match the slow decay of GCaMP.

#### Lick choice

Two binary regressors (ipsilateral, contralateral), modeled as unit impulses at each trial’s lick time, also convolved with the calcium kernel.

#### Movement

Seven finite-impulse-response (FIR) lag regressors derived from whole facial motion energy (ME), spanning lags of −0.1 to +0.5 s in 0.1 s steps.

#### Pupil/Arousal

Seven FIR lag regressors from pupil diameter, spanning 0 to +0.6 s in 0.1 s steps. All regressors were z-scored column-wise prior to fitting, placing sparse event regressors (sound, lick) and dense continuous regressors (movement, pupil) on a comparable scale for regularization.

Model fitting. We used ridge regression with a data-adaptive regularization parameter λ = α · mean(diag(XᵀX)), where α = 0.1 and the diagonal was computed over non-intercept columns only; the intercept was not penalized. Model performance was assessed by 5-fold cross-validation with fold assignment at the trial level (all timepoints within a trial assigned to the same fold), preventing leakage due to temporal autocorrelation within trials. Cross-validated R² was computed on held-out data as:

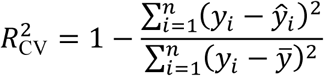

Variance decomposition. Unique variance attributable to each predictor group was estimated as ΔR² = R²_full_ − R²_without_predictor_, clamped to zero if negative. Shared variance was defined as R²_full_ − Σ(ΔR²_unique_). Cells were included if R²_full_ > 0. (n = 1390 of 1479 tracked cells across all mice).

### Statistics

All statistical analyses were run in MATLAB/python. Significance levels *, **, and *** correspond to p-values lower than 0.05, 0.01, and 0.001, respectively. All descriptive values are mean and standard deviation unless otherwise noted.

For comparisons across behavioral conditions (e.g., active vs. passive, or trial-type TMI), statistical significance was assessed using a linear mixed-effects model with mouse as a random effect (val ∼ cond + (1|mouse)), controlling for inter-animal pseudo-replication while treating cells as independent observations within each mouse. For silencing comparisons, where each cell contributed paired control and silenced observations, a fully nested model (y ∼ Cond + (1|Mouse) + (1|Cell)) was used, removing between-cell variance and providing a paired-test equivalent within a mixed-effects framework. BH-FDR corrected across all tests within each analysis.

## Results

### Mice learned the head-fixed 2AFC licking task at variable rates

The task in this study was a two alternative forced choice task under head-fixation using licking as the choice (Fig. 1A). The mice were first trained in a training room before being trained under the 2P microscope. The two pure tones to differentiate are 8 kHz and 16 kHz. Fig. 1B presents the learning curves of individual mice along the training days starting after the habituation days. This figure shows that different mice learned the same task with different rates, with some learning it within 5 days while others needed more than 2 weeks.

A camera from the front angle of the mouse was used to record the face and tongue movement (Fig. 1C) which produced the videos to study the pupil and movement of the mice.

### Mice reliably discriminated 8 and 16 kHz tones during 2P imaging

Seven mice successfully learned the frequency discrimination task and went to the 2P imaging phase. Mean accuracy of the behavioral performance under 2P microscope was 70.0 ± 9.6% for 8 kHz trials and 69.7 ± 8.5% for 16 kHz trials (Fig. 2A). Several mice performed slightly below the 75% criterion during imaging sessions. We attribute this to the additional stress of active 2P imaging, despite prior habituation and training under the microscope without active scanning. Choice response latencies were significantly shorter for 8 kHz hit trials (0.62 ± 0.01 s) than for 16 kHz hit trials (0.90 ± 0.09 s; paired t-test, p = 0.028; Fig. 2B, C).

**Fig. 2.**
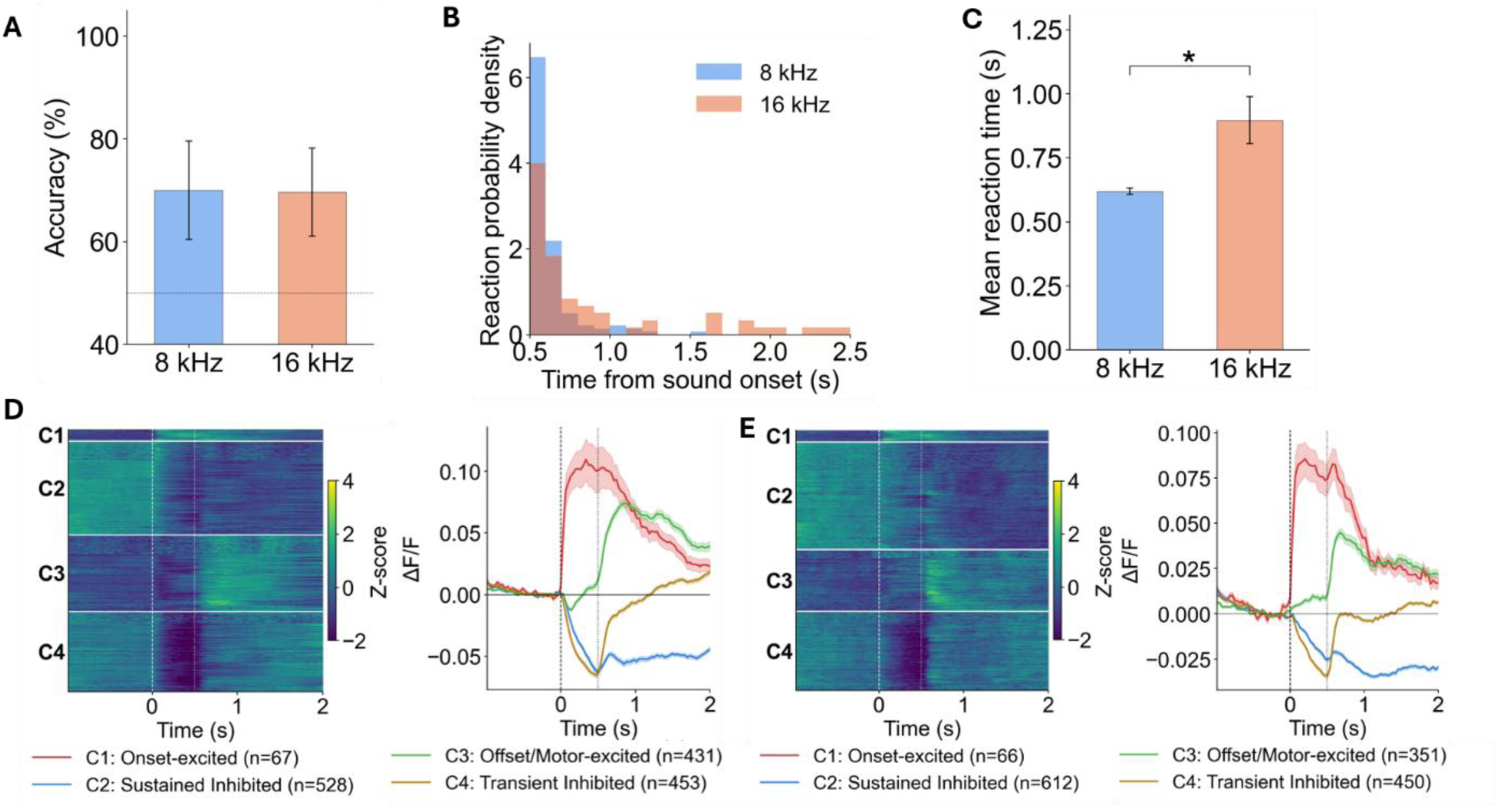
Behavioral performance and DCIC response classes during the 2AFC tone discrimination task. (A) Behavioral accuracy for 8 kHz and 16 kHz target tones. The dotted horizontal line indicates chance performance at 50%. (B) Distribution of reaction times for 8 kHz and 16 kHz trials, measured as the time from sound onset to the first decision lick. (C) Mean reaction time for 8 kHz and 16 kHz trials. Mice responded significantly more slowly to 16 kHz trials than to 8 kHz trials. (D–E) Sound-evoked DCIC population responses to (D) 8 kHz and (E) 16 kHz tones. Left panels show z-scored ΔF/F responses of individual neurons, grouped into four response classes according to their temporal response profiles. Rows represent individual neurons, and the vertical dashed line indicates sound onset. Right panels show the mean ΔF/F traces for each response class. C1 neurons showed onset-excited responses, C2 neurons showed sustained inhibited responses, C3 neurons showed offset/motor-excited responses, and C4 neurons showed transient inhibited responses. Numbers in parentheses indicate the number of neurons in each class. Shaded regions indicate SEM. Bars represent mean ± SEM. *p < 0.05.

### DCIC neurons formed consistent functional response clusters across 8 and 16 kHz trials

Clustering of 8 kHz active-trial PSTHs identified four functional groups (Fig. 2D): Onset-excited (C1, n = 67), characterized by a sharp tone-onset response; Sustained Inhibited (C2, n = 528), showing prolonged suppression throughout the trial; Offset/Motor-excited (C3, n = 431), with a delayed response peaking at the post-sound window (∼0.5–1.5 s); and Transient Inhibited (C4, n = 453), exhibiting brief inhibition at tone onset followed by recovery. Applying the identical pipeline to 16 kHz active-trial PSTHs from the same 1,479 neurons yielded four clusters (Fig. 2E): Onset-excited (C1, n = 66), Sustained Inhibited (C2, n = 612), Offset/Motor-excited (C3, n = 351), and Transient Inhibited (C4, n = 450). These results demonstrate a high degree of consistency in the functional classification of neurons across different trial types (8 kHz vs. 16 kHz), as evidenced by the emergence of nearly identical response profiles with comparable cluster sizes. The functional clustering was consistent across 8 kHz and 16 kHz trials (nearly identical response profiles and comparable cluster sizes). For clarity of presentation, subsequent analyses focus on 8 kHz trials.

### Active listening modulated DCIC response activity

We studied the neural activity in trials of 8 kHz pure tone in both active phase and passive phase (Fig. 3A). The Onset-excited cluster (C1) showed no significant active vs. passive difference in either window. The Sustained Inhibited cluster (C2) was significantly more suppressed during active trials in the post-sound window (p < 0.001), but not in the sound window. The Offset/Motor-excited cluster (C3) showed significantly greater active than passive responses in both windows (sound: p < 0.001; post-sound: p < 0.001). The Transient Inhibited cluster (C4) was more suppressed during active trials in both windows (sound: p < 0.001; post-sound: p < 0.001).

**Fig. 3.**
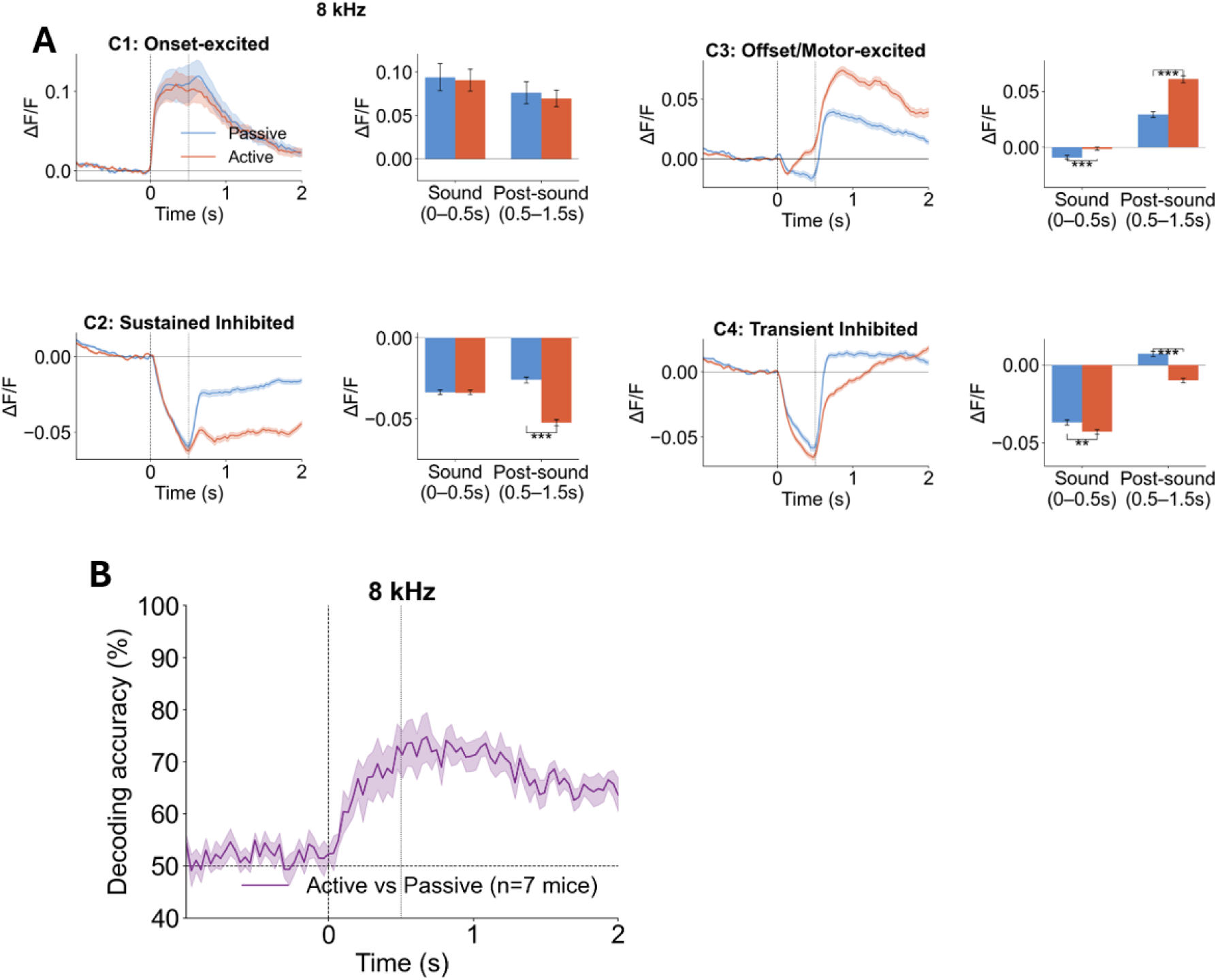
Task engagement modulates DCIC responses to 8 kHz tones. (A) Mean calcium responses of four DCIC response clusters to 8 kHz tones during passive listening and active task performance. For each response class, left panels show mean ΔF/F traces in passive trials (blue) and active trials (orange), and right panels show the average response amplitudes during the sound window (0–0.5 s from sound onset) and post-sound window (0.5–1.5 s from sound onset). The vertical dashed line indicates sound onset, and the vertical gray line indicates sound offset. C1 neurons were onset-excited, C2 neurons were sustained inhibited, C3 neurons were offset/motor-excited, and C4 neurons were transient inhibited. Task engagement altered response amplitudes in a cluster- and time-window-dependent manner, with prominent active-passive differences in the post-sound responses of C2, C3, and C4 neurons. C1: n=67; C2: n=528; C3: n=431; C4: n=453. (B) Time-resolved SVM decoding of behavioral state from DCIC population activity in response to 8 kHz tones. A support vector machine classifier was used to discriminate active-task trials from passive-listening trials based on neural activity at each time point. The decoding accuracy rose above chance after sound onset, indicating that DCIC population activity contained information about behavioral state during and after the sound response. The dashed horizontal line indicates chance-level decoding at 50%. The vertical dashed line indicates sound onset, and the vertical gray line indicates sound offset. Shaded regions indicate SEM across mice. Bars and traces represent mean ± SEM. Asterisks indicate significant differences between active and passive conditions. n = 7 mice.

We used a SVM model to test whether the populational neural activity encoded the context of active or passive listening. Context decoding of the 8-kHz tone trials rose sharply above chance (50%) at sound onset and peaked at approximately 70–75% accuracy during the sound window (0–0.5 s), remaining significantly above chance throughout the post-sound window (0.5–1.5 s) before gradually returning toward baseline (Fig. 3B). This sustained above-chance decoding indicates that DCIC population activity encodes task context in a manner that outlasts the acoustic stimulus itself.

### Hit and miss trials drove outcome-dependent modulation of DCIC activity

The active condition pools heterogeneous behavioral outcomes, and the magnitude of the combined active-versus-passive effect is therefore influenced by the relative proportion of hit, miss, and no-lick trials. To localize the modulation to specific behavioral states, we further separated the active trials into hit, miss and no-lick trials according to the behavioral outcome. The four clusters showed distinct and outcome-dependent patterns of modulation across trial types (Fig. 4). C1 neurons showed no significant difference between passive and any active trial type in either window, indicating that onset responses are driven by the auditory stimulus and are insensitive to behavioral outcome. C2 neurons under the post-sound window were more strongly suppressed during hit (p <0.001) and miss (p <0.001) trials than during passive trials, with no-lick trials resembling the neural activity of the passive trials. C3 neurons were more strongly excited during hit and miss trials than during passive trials (both p < 0.001), with passive trials being more strongly excited than the no-lick trials (p<0.001). C4 neurons showed deeper inhibition in the post-sound window during hit trials relative to passive (p < 0.001). These results indicate that the major task modulation was driven by hit and miss trials and the no-lick trials might have an opposite minor effect of task modulation.

**Fig. 4.**
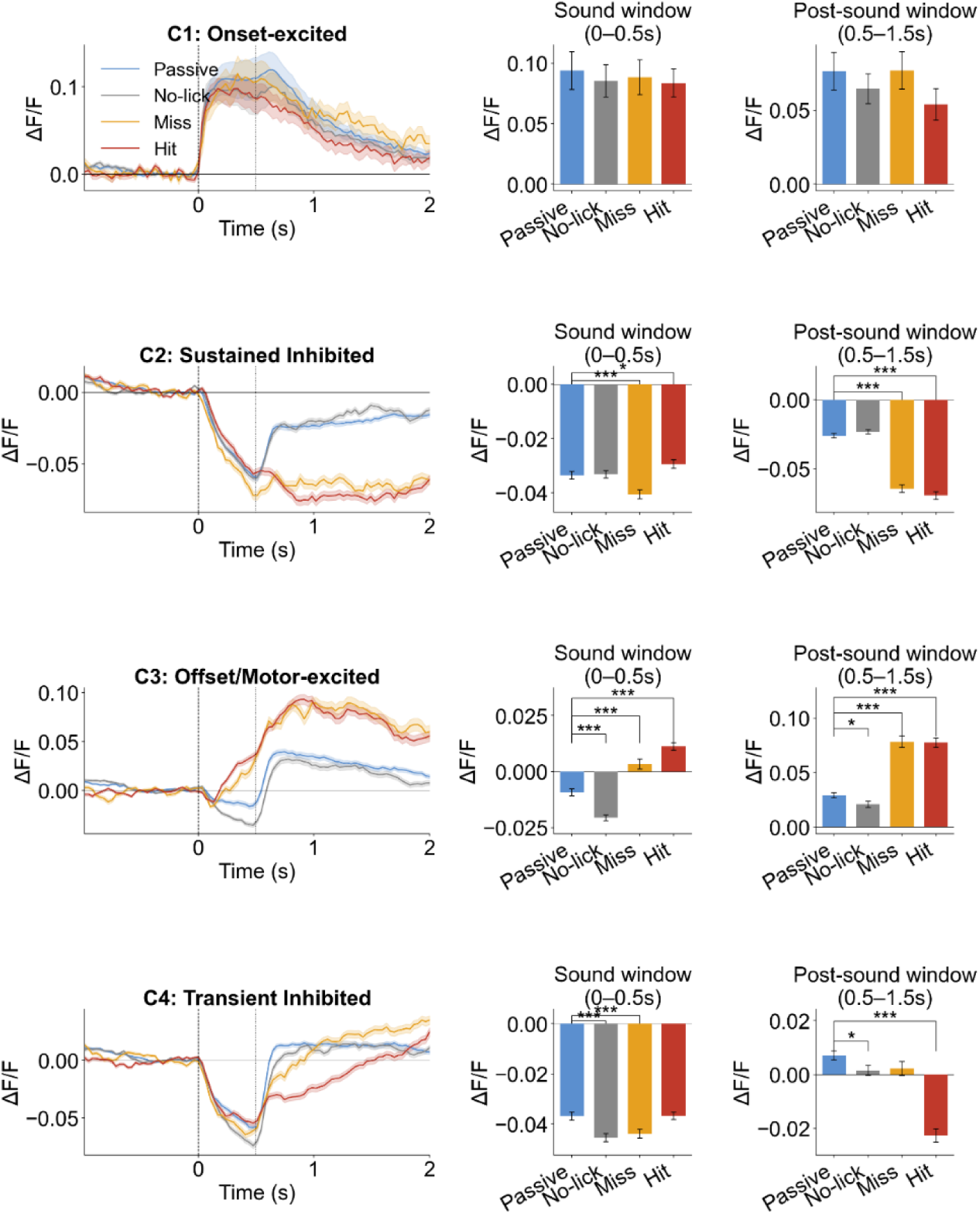
Outcome-dependent modulation of 8 kHz responses in DCIC neurons during task performance. Mean calcium responses to 8 kHz tones were compared between passive listening and active-task trials separated by behavioral outcome. For each response class, the left panel shows the mean ΔF/F trace for passive trials, no-lick trials, miss trials, and hit trials. The middle and right panels show the average response amplitudes during the sound window, defined as 0–0.5 s from sound onset, and the post-sound window, defined as 0.5–1.5 s from sound onset, respectively. The vertical dashed line indicates sound onset, and the vertical gray line indicates sound offset. C1 neurons showed onset-excited responses, C2 neurons showed sustained inhibited responses, C3 neurons showed offset/motor-excited responses, and C4 neurons showed transient inhibited responses. Behavioral outcome modulated DCIC activity in a response-class-dependent manner. C1 responses showed relatively weak outcome dependence, whereas C2, C3, and C4 neurons showed stronger differences between passive, no-lick, miss, and hit trials, particularly during the post-sound window. C1: n=67; C2: n=528; C3: n=431; C4: n=453. Traces and bars represent mean ± SEM. Asterisks indicate significant pairwise comparisons between trial types. *p < 0.05, ***p < 0.001, ****p < 0.0001.

### Functional response clusters showed distinct sensory and motor encoding profiles

To quantify the contribution of sensory, motor, and arousal signals to the activity of each functional cluster, we fit a linear encoding model to each cell’s ΔF/F using four behavioral predictors: sound identity, lick choice direction, facial movement energy, and pupil size. The four clusters differed markedly in which predictors drove their activity and how much the activity was driven by these predictors (Fig. 5A). C1 neurons had the highest total explained variance (mean R² = 0.161) and were dominated by sound: the sound predictor accounted for 77.7% of total R², with only minor contributions from movement (8.1%), lick choice (3.5%), and pupil/arousal (2.1%). This profile is consistent with C1 representing classical auditory-responsive neurons tightly locked to sound onset. C2 neurons showed a more distributed variance profile (mean R² = 0.087), with roughly equal contributions from sound (28.1%), movement (27.7%), and a substantial shared component (31.7%), indicating that C2 activity is co-modulated by both auditory and motor signals. C3 neurons (mean R² = 0.088) were most strongly explained by movement (36.8%), followed by lick choice (15.6%), while sound accounted for only 13.4%. C4 neurons had the lowest total explained variance (mean R² = 0.059) and were moderately sound-driven (42.1%), with secondary contributions from movement (27.6%). Pupil/arousal contributed minimally across all clusters (2–6% of total R²).

**Fig. 5.**
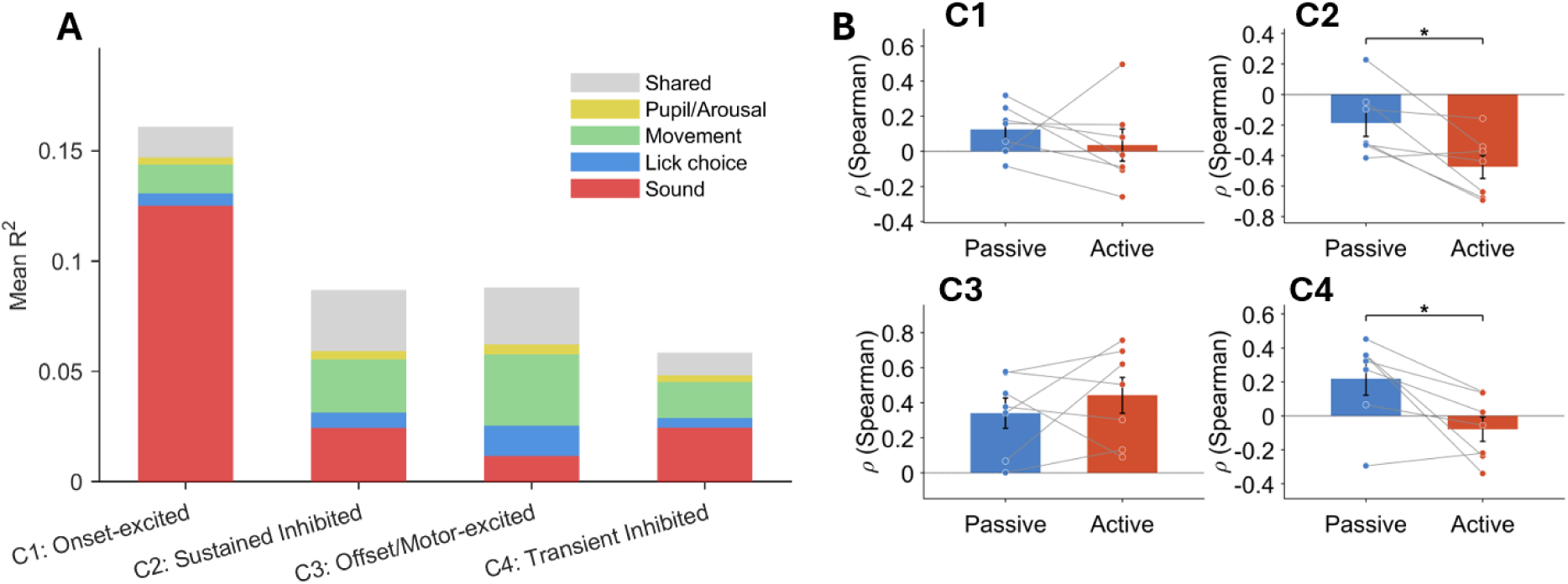
Encoding model analysis and task-dependent movement–activity coupling across DCIC functional clusters. (A) Mean explained variance decomposition from a linear encoding model fit to each neuron’s ΔF/F activity during active-task trials. The model included predictor groups for sound identity, lick choice, facial movement energy, and pupil/arousal. Stacked bars show the mean unique contribution of each predictor group to the full-model R² for each functional cluster, together with the shared component of explained variance. Shared variance indicates the portion of the full-model R² that was not uniquely attributable to a single predictor group. (B) Task-dependent coupling between mouth motion energy and DCIC activity. Mouth motion energy was measured from a video ROI around the mouse’s mouth and used as a proxy for licking and orofacial movement. For each mouse, Spearman correlations were computed between mouth motion energy and trial-by-trial neural activity during the post-sound window, defined as 0.5–1.5 s after sound onset, separately for passive-listening and active-task trials. Bars show the mean correlation across mice, and gray lines connect paired passive and active values from the same mouse. Bars represent mean ± SEM. *p < 0.05. n = 7 mice.

### Movement–activity coupling was context-dependent in inhibited DCIC clusters

Because the DCIC was shown to contain a large percentage of motion-modulated neurons (Han et al., 2026), and our encoding model showed that movement contributed a substantial proportion of the explained variance in C2, C3 and C4, we asked a further question that the encoding analysis could not address: whether the relationship between movement and neural activity was itself dependent on task context. The encoding model was fit on active trials only and therefore quantified how much movement contributed, but not whether this movement–activity coupling differed between active and passive listening. We therefore compared the correlation between movement and neural activity across the two contexts.

Because the activity that distinguished hit and miss trials from no-lick trials coincided with licking, we reasoned that the movement most relevant to this coupling was orofacial/licking movement. As licking was only registered at the choice tube and could not be tracked during passive trials, we marked a ROI around the mouse’s mouth in the video and computed the frame-to-frame motion energy (mouth ME) as a proxy for licking and orofacial movement. We then computed the Spearman correlation between mouth ME and the trial-by-trial activity of C1–C4 neurons during the post-sound window (0.5–1.5 s), separately for active and passive trials, and compared the two (Fig. 5B).

The coupling between mouth ME and neural activity differed between contexts in a cluster-specific manner. In C1 (Onset-excited), the correlation was weakly positive and did not differ between active and passive trials. In C3 (Offset/Motor-excited), mouth ME was strongly positively correlated with activity in both contexts, with no significant difference between them (passive ρ = 0.34, active ρ = 0.44; p = 0.350), indicating a context-independent motor coupling. In contrast, both C2 and C4 showed a significant context-dependent shift. In C2 (Sustained Inhibited), the negative correlation with mouth ME was significantly stronger during active than passive trials (mean Δρ = −0.31; paired t, p = 0.021; 6/7 mice). In C4 (Transient Inhibited), the correlation reversed sign, from positive during passive trials to near-zero or negative during active trials (mean Δρ = −0.28; p = 0.026; 6/7 mice), although inter-animal variability was substantial.

### Auditory cortex silencing selectively altered task modulation in Sustained inhibited DCIC neurons

The analyses above establish that DCIC activity is modulated by task context. We finally asked whether this modulation depends on descending input from the auditory cortex, a major source of corticocollicular projections to the DCIC (Bajo & King, 2012). We bilaterally silenced AC with CNO in three hM4D(Gi)-bilateral-AC-expressing mice. We performed behavior and imaging on separate days for the control (saline) and AC-silencing conditions, with two mice receiving the control day first and one receiving the CNO day first. The task performance was not significantly affected by the AC silencing, although there was a normal day-to-day accuracy variance (Fig. 6A). In all the recorded neurons in the three mice, we found 171 matched neurons which we could identify on both days.

**Fig. 6.**
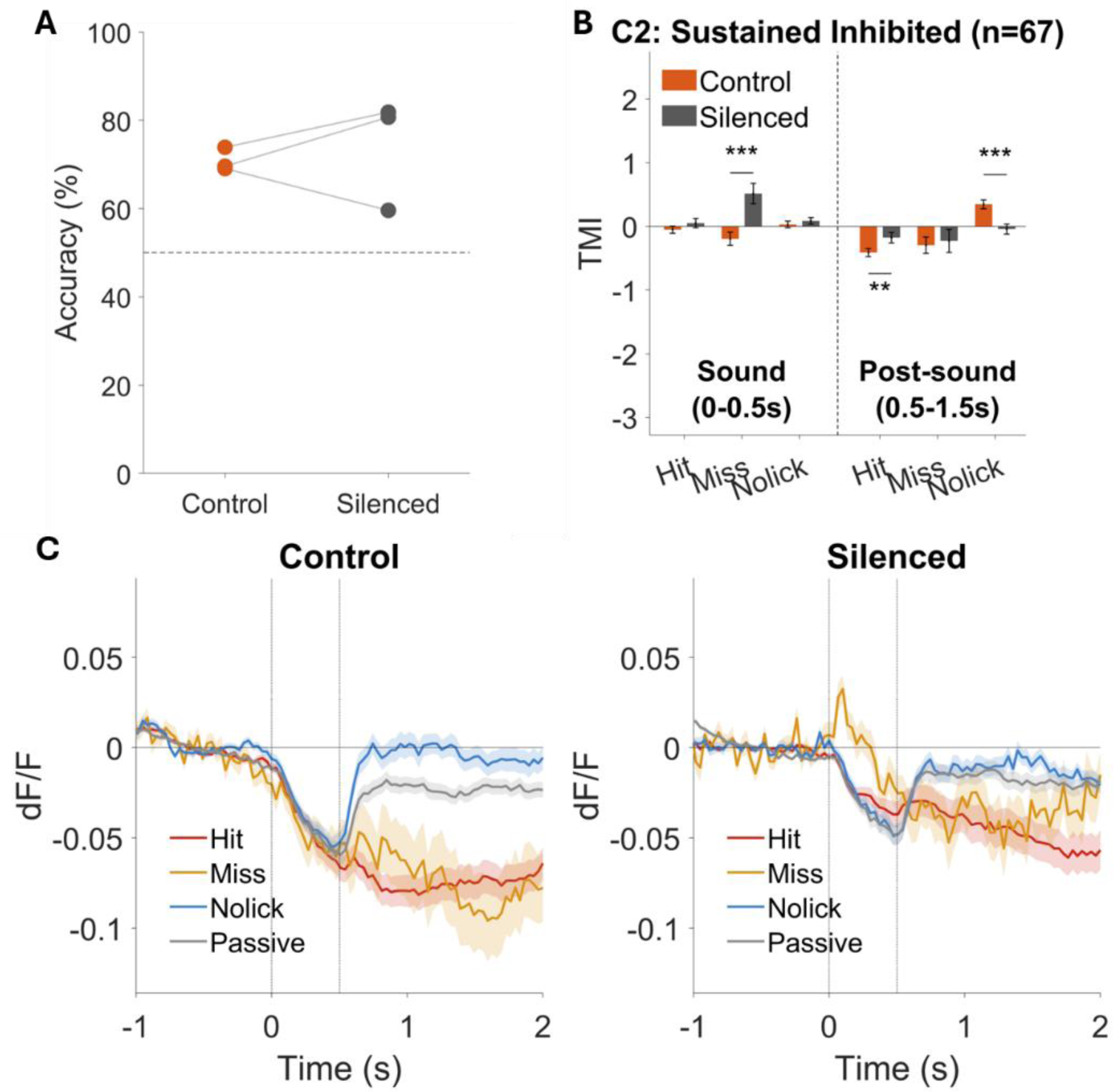
Auditory cortex silencing and task modulation of C2 DCIC neurons. (A) Behavioral accuracy during control and auditory cortex-silenced sessions. Control sessions were performed after saline injection, and silenced sessions were performed after CNO injection in hM4D(Gi)-expressing mice with bilateral auditory cortex targeting. Each dot represents one mouse, and gray lines connect paired control and silenced sessions from the same mouse. The dashed horizontal line indicates chance performance at 50%. (B) Task modulation index (TMI) of C2 sustained-inhibited neurons under control and auditory cortex-silenced conditions. TMI was calculated as (*active* − *passive*)/(∣ *active* ∣ +∣ *passive* ∣), where active responses were measured separately for hit, miss, and no-lick trials, and passive responses were measured during passive listening. TMI values were computed during the sound window, defined as 0–0.5 s after sound onset, and the post-sound window, defined as 0.5–1.5 s after sound onset. The comparison was done between matched C2 neurons identified across control and silenced sessions. (C) Mean ΔF/F traces of C2 sustained-inhibited neurons during control and auditory cortex-silenced sessions. Traces show responses during hit, miss, no-lick, and passive trials. Vertical gray lines indicate sound onset and sound offset. Shaded regions indicate SEM. For all panels, control indicates saline sessions and silenced indicates CNO sessions. **p < 0.01, ***p < 0.001. n = 3 mice; 171 matched neurons across sessions; C2 neurons, n = 67.

We quantified task modulation using the task modulation index (TMI = (active − passive) / (|active| + |passive|)), where positive values indicate stronger active-state responses and negative values indicate stronger passive-state responses. AC silencing significantly altered the TMI of only the C2 (Sustained inhibited) neurons (Fig. 6B) in which the TMI of miss trials became more positive in the sound window and the TMI of hit trials became more positive and the TMI of the no-lick trials became more negative in the post-sound window. The neural activity traces showed that the active-trial traces in the post-sound window became closer to the passive trials under the silenced condition (Fig. 6C) which indicates that the AC silencing might decrease the task modulation in C2.

## Discussion

Our data show that DCIC neural activity is modulated by task context, that this modulation is cell-type-specific across functionally defined clusters, and that the modulation of one cluster — the Sustained Inhibited population (C2) — depends on auditory cortex activity.

### Task modulation is closely tied to movement

A central question is whether this modulation reflects a cognitive process such as task engagement, or movement accompanying the task. Our encoding model showed that movement contributed a substantial fraction of the variance in C2, C3 and C4. Moreover, the direction of the movement–activity correlation matched the direction of the task modulation itself: in C2, which was suppressed during active trials, activity was negatively correlated with mouth motion energy. On top of this baseline coupling, the correlation was further strengthened during active relative to passive trials in C2 and C4. Meanwhile, C3 activity was positively correlated with mouth motion in both contexts, and this coupling did not differ between active and passive trials. This positive correlation is consistent with the stronger C3 excitation on hit and miss trials, where movement is greater: C3 modulation is well explained by a context-independent coupling to movement, identifying it as a predominantly movement-driven cluster. Because mouth motion is dominated by licking, much of the task modulation — both its direction and its enhancement during active trials — is likely movement-related rather than reflecting a movement-independent cognitive signal. This is consistent with Han et al. (2026), who found that locomotion drives IC activity even in the absence of auditory input, indicating that the IC receives movement-related signals independent of sound — as we observe for the movement-coupled task modulation here. We are therefore cautious in interpreting it as task engagement per se: we found no evidence that the active–passive modulation is independent of movement. Two possibilities remain and cannot be distinguished here — that task context genuinely modulates how movement is coupled to DCIC activity, or that the movements themselves differ (licking at the choice tube versus spontaneous licking during passive trials). Distinguishing them will require trial-by-trial detection of licking, including spontaneous licks during passive trials, which our tube-based readout cannot capture; video-based pose estimation is an important future direction.

### Auditory cortex contributes to the modulation of C2

Silencing AC selectively altered the task modulation of C2, leaving the other clusters unaffected. Because we silenced AC broadly rather than a defined projection class, we cannot attribute this to a specific pathway; what our data establish is that AC activity is required for the normal task modulation of C2 alone. Corticocollicular projections are glutamatergic and excitatory at their terminals (Suga, 2020), so silencing them, which reduces an inhibitory form of modulation, suggests the cortical influence on C2 is exerted indirectly, through local inhibitory circuitry within the IC. The insensitivity of the other clusters indicates their modulation does not require AC and must be supported by other inputs — other descending sources, neuromodulatory systems (Ayala & Malmierca, 2015; Kuchibhotla et al., 2017), or movement-related signals reaching the IC independently of AC. Our data thus point to at least two dissociable influences on behavioral-state modulation in the DCIC.

### Limitations

Our silencing experiment lacked a CNO-only control in DREADD-negative animals; because systemic CNO can have off-target effects via back-metabolism to clozapine (Martinez et al., 2019), we cannot fully exclude a CNO contribution, though a non-specific effect would be expected to act broadly rather than confined to C2. This control, and replication in a larger cohort (here three mice, 171 matched neurons), remain for future work.

## Acknowledgements

This work was supported by NIH NIDCD Grants R01DC016599, R01DC013073, and R21DC021605. We thank Yuan Cai, Kush Nandani, Grace Ledogar, Austin Douglas, Vraj Thakkar, Michael Chen for training the mice.

